# Honey bees solve a multi-comparison ranking task by probability matching

**DOI:** 10.1101/2020.03.02.967661

**Authors:** HaDi MaBouDi, James A.R. Marshall, Andrew B. Barron

## Abstract

Honey bees forage on a range of flowers, all of which can vary unpredictably in the amount and type of rewards they offer. In this environment bees are challenged with maximising the resources they gather for their colony. That bees are effective foragers is clear, but how bees solve this type of complex multi-choice task is unknown. Here we challenged bees with a five-comparison choice task in which five colours differed in their probability of offering reward and punishment. The colours were ranked such that high ranked colours were more likely to offer reward, and the ranking was unambiguous. Bees choices in unrewarded tests matched their individual experiences of reward and punishment of each colour, indicating bees solved this test not by comparing or ranking colours but by matching their preferences to their history of reinforcement for each colour. We used a computational model to explore the feasibility of this probability matching strategy for the honey bee brain. The model suggested a structure like the honey bee mushroom body with reinforcement-related plasticity at both input and output was sufficient for this cognitive strategy. We discuss how probability matching enables effective choices to be made without a need to compare any stimuli directly, and the utility and limitations of this simple cognitive strategy for foraging animals.

## Introduction

A honey bee’s foraging ecology presents a prodigious cognitive challenge. Bees must gather concealed pollen and nectar from many cryptic and variable flower species, all of which vary moment by moment in the quality and amount of reward. None of these probabilities are known to the foraging bee. This kind of problem has been described as a multi-armed bandit task [1, 2]: a task in which there are multiple options available all with unknown potential for variable payouts. Here we challenged honey bees with a controlled learning task that offered five options differing in the probability of offering reward and punishment to examine what bees learned and how they solved this type of multiple-comparison task.

One solution to this kind of task would be to identify the option that offered the highest probability of reward and always pick that option (probability maximising)[3]. For example, if a bee learned during training that blue is rewarded 60% of the time and yellow is rewarded 40% of the time, and if the bee is able to compare and rank these two alternatives then she would maximise her return by picking the blue option all the time. An alternative solution to this type of task involves no comparisons at all. Bees could simply learn the reinforcement properties of each option and match their choices accordingly (probability matching) [4–6]. In the above example, post training bees would choose blue 60% of the time and yellow 40% of the time. This could be achieved simply by learning the reinforcement history of each stimulus without any comparison or ranking of them.

Probability maximising is the economically rational solution since it offers maximum possible reward in the kind of scenario described above where probabilities of different options are known and fixed. But in scenarios in which the reward probabilities are unknown and/or changing probability matching offers a better solution to the trade-off of exploiting current resources while exploring other alternatives [2, 7]. Hence in many ecological situations probability matching may be an evolved optimal strategy [1, 2, 7, 8].

Probability maximising requires a capacity to compare the value of different options and to rank them accordingly. Until recently this kind of cognition was presumed to be too complex for insects, but bees do have the capacity to learn abstract relationships between stimuli [9–11], and a recent study with Polistes wasps showed the wasps could pass a behavioural test considered indicative of the capacity for transitive inference [12]. This requires arranging stimuli in a ranked order [12] suggesting that *Polistes dominula* and *Polistes metricus* at least have a capacity for ranking of stimuli.

Probability maximising requires different options be compared. By contrast probability matching involves no comparisons: it simply involves learning the reinforcement properties of each stimulus. This has been considered less intelligent than probability maximising by some [4, 13]. While that interpretation is open to debate [humans often probability match when making choices [14], it is true that probability matching is computationally simpler than probability maximising [2, 7].

Both honey bees and bumblebees are clearly able to quickly learn features of stimuli that offer rewards, and adapt their behaviour when reward contingencies change [2, 15-17]. Whether honey bees choose by learning the probability of reinforcement for different stimuli (probability matching), or whether they rank stimuli by probability of reinforcement (probability maximising) is presently unclear.

Greggers and Menzel [16] trained honey bees to four different artificial feeders that varied in rate of sucrose delivery, and in the same study bees showed evidence of both probability matching and probability maximising strategies [16]. Fischer et al [17] and Keasar et al [2] set honey bees and bumblebees *(Bombus terrestris)* and found that bees imperfectly matched their preferences to feeder probability of reward. In these earlier studies it is possible that the binary choice paradigm (equivalent to a two-armed bandit task) was not complex enough for bees to use a probability maximising strategy. Probability matching is a very common choice strategy for humans, but we will switch to probability maximisation if the ‘stakes’ or complexity of the choice are increased [18]. Adding punishment to a choice test increases the negative consequence of a wrong choice for honey bees, which enhances discriminant colour learning and reduce generalisation [19]. Here we examined honey bees’ choice strategy in a complex multi-option choice test in which bees were punished for wrong choices.

We set honey bees a five-armed bandit task. We challenged honey bees with a colour learning task in which five coloured stimuli were organised in a rank order such that when trained with pairwise presentations of colours only the higher ranked colour was rewarded, with the lower ranked colour being punished. The ranking was unambiguous, and it influenced the probability of each colour being rewarded or punished during training. Tests then explored whether honey bees demonstrated learning of the ranking of the colours, or whether they instead learned the reinforcement history of each separate colour. Our data show honey bees rapidly learned to match their choices in tests to the reinforcement history of each colour. We developed a neural model to explore the feasibility of probability matching in a five-arm bandit task for an insect brain.

## Material and methods

### Animals and testing arena

Experiments were conducted at the Sheffield University Research Apiary. The apiary contained four standard commercial hives of honey bees (*Apis mellifera*). To attract honey bees for our experiments we placed a gravity feeder containing 20% sucrose solution (w/w) approximately 15m from the hives. Bees visiting the gravity feeder were given individually distinctive marks with coloured paints on their abdomen and/or thorax using coloured Posca marking pens (Uni-Ball, Japan).

5 m from the gravity feeder (and further from the hives) we established the testing arena. This was a box (100 x 80 x 80 cm) made from white expanded PVC foam boards. The roof was UV-transparent Plexiglas. Bees could enter the arena through a transparent Perspex corridor (20 x 4 x 4 cm). Interior walls and the ground were covered with a pink random dot pattern to provide a contrast between the colour of the bees and the background to assist further video analysis (Fig. 1A).

**Figure 1.**
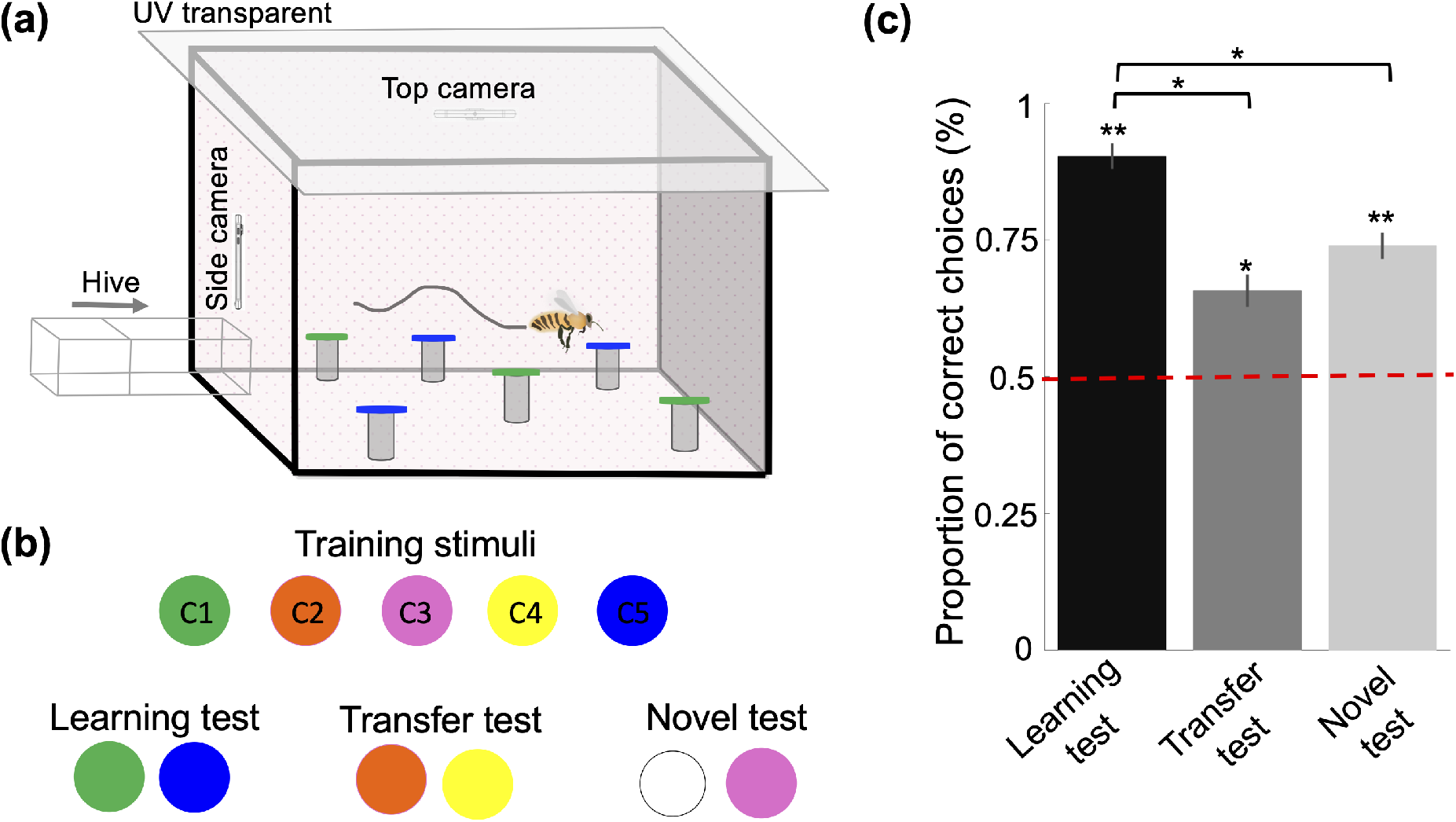
***a*)** The experimental paradigm. Each bee received 18 training trials. In each training trial, a free-flying marked bee was presented with eight stimuli, four of each of two colours. Stimuli of one colour offered 10 ul sucrose solution 50% (w/w) and the other colour offered 10 ul of saturated quinine hemisulphate. ***b)*** Five different colours were used in this study. The colours differed in the proportion of training trials in which they offered reward and punishment (rewarded at 100%, 66%, 50%, 33%, 0% of trials, Table S1). Following training bees received three unreinforced tests. In the *learning test* bees were presented with the colours that had always been rewarded and always punished in training. In the *transfer test* bees were tested with a colour combination that had not been used in training and that differed in their reward frequency in training trials (66% trials rewarded vs 33% of trials rewarded). In the *novel test* bees were presented with a novel colour and a colour that had been rewarded in 50% of training trials. ***c)*** Performance on the tests. Bars show mean frequency with which bees landed on stimuli they had experienced as more rewarded during training. In the novel test, bees chose the trained colours (C3) more than the novel colour. N = 20. Vertical lines: s.e.m. Dashed line indicates performance expected at random. * indicates p<0.05 and ** indicates p<0.005

### Stimuli

Bee were trained to visit coloured stimuli inside the arena. These were disks (2.5 cm in diameter) of coloured paper covered with transparent laminate (Figure 1 and S1). Stimuli were placed on small inverted transparent plastic cups in the arena to raise them from the arena floor.

### Pre-training phase

Marked bees were attracted away from the gravity feeder by offering them a cotton bud soaked with 50% sucrose solution (w/w). Once a bee began feeding from the cotton bud, she was gently moved to the entrance of the flight arena and there provided more 50% sucrose to drink to satiation. This procedure was repeated until the bee flew independently to the entrance of the arena. Bees were trained to fly into the flight arena via the entrance tube to find drops of 50% sucrose placed on transparent disks of laminate on top of the plastic cups. Bees were released from the arena by lifting the roof. Once a bee flew by herself into the arena to feed, she was selected for the training phase.

### Task and training

Bees were trained with five different coloured stimuli in a colour discrimination task. The five different colours were assigned an arbitrary number (C1-5). In each trial a bee was presented with eight stimuli: four of one colour and four of a different colour. For any given pair of colours presented in a trial the colour with the lower number was rewarded and the colour with the higher number was punished, therefore whether a colour was rewarded or punished in training depended on its number and the number of the colour it was paired with (Table S1). Feeders were placed randomly within the arena. Feeders were rewarded with 10 μl sucrose solution 50% (w/w) and punished with 10 μl of saturated quinine hemisulphate solution.

In each trial bees were able to freely land on stimuli and to feed from rewarded stimuli. 10 μl drops of 50% sucrose solution were replaced on depleted rewarded stimuli until the bee had fed to satiation and left the arena via the roof. Bees returned to the arena by their own volition. Typically, the inter-trial interval was 5-10 minutes. After each training trial, all stimuli were cleaned with 70% ethanol to remove any possible pheromonal cues left by the bee.

Bees were assigned at random to one of two groups: A and B. For group A colours were assigned as: blue = C1, yellow = C2, pink = C3, orange = C4 and green = C5 with white used as a novel colour for testing. For group B the colour assignment was: green = C1, orange = C2, white = C3, yellow = C4 and blue = C5 with pink used as a novel colour in testing. Comparing behaviour of bees from groups A and B allowed us to explore for possible innate colour biases influencing choice.

Over 18 training trials bees experienced all combinations of the five colours twice, with the exception that bees in training never experienced C2 paired with C4 (Table S1& S2). This pairing was excluded from training so that in the post training *transfer test* we could examine how trained bees responded to a colour pair they had never previously encountered. The 18 bouts of training presented bees with reward from C1 in eight out of eight trials; from C2 in 4 out of 6 trials, from C3 in 4 out of eight trials, from C4 for two out of six trials, and never from C5 (Table 1).

We established two different sequences for presentation of training trials: protocols P1 and P2 (Table S2). Bees were randomly assigned to a protocol and comparing performance between the two protocols allowed us to examine whether training sequence influenced performance. In training each bee was therefore assigned to one colour group (A or B) and one training protocol (P1 or P2).

### Testing

Immediately following the training phase each bee was given three learning tests. The *learning test* presented bees with C1 and C5 – a colour combination they had previously experienced. The *transfer test* presented bees with C2 and C4 – a combination they never experienced in training. The *novel test* presented bees with white and pink, which was a choice between a colour they had experienced to be rewarded and punished equally often and a colour they had not experienced in training. During all tests all stimuli offered 10 ul water. During tests bees were observed and video recorded for 120 s and their landings on stimuli recorded, after which bees were released from the roof of the arena. The sequence of the three tests was randomised for each bee. Between tests bees were allowed to feed in the arena on 10 ul sucrose drops placed on disks of transparent laminate so that we maintained their motivation to visit the arena. As in training, stimuli were cleaned between each test.

### Observation and video recording

To record bee behaviour inside the arena iPhone 6 cameras were positioned above the entrance viewing into the arena, and on the top of the arena viewing down. The camera was configured to record at 30 fps at a resolution of 720p (1,280x 720 pixels) in the training phase, and 240 fps in the testing phase. During training we recorded from when the bee entered the arena until she was released from the roof. During tests we recorded 120 s from when she entered the arena. A sample recorded video is provided as supplementary Videos S1. We developed an algorithm to analyse the flight paths of bees from the videos. Bees’ flight path was determined by extracting the x-y coordinates of the bee’s body and their direction of flight frame by frame (Video S1).

Only bees that completed the entire training and test sequences were included in the results. Of our 20 bees, only one bee did not complete the entire paradigm due to rain stopping the experiment on that day We noted each time a bee landed on a stimulus in training, from which we determined how many times each bee encountered each colour as rewarded or punished.

## Data analysis and statistics

To evaluate bees’ performance in the three different unrewarded tests, we analysed the proportion of correct choices estimated as landings on the rewarded stimuli divided by the total number of landings during the 120 second test. We examined the effect of colour group (A or B), protocol (P1 or P2), test type and colour ranking on performance using Wilcoxon signed rank and Wilcoxon rank-sum tests.

To further examine whether individual experience of the history of reward and punishment associated with each colour influenced degree of colour preference we calculated the number of visits to each colour that was rewarded and punished for each bee (Figure S3). In our protocol bees freely visited and chose multiple feeders within each training bout and therefore each individual bee experienced a unique history of rewards and punishments for each colour.

For each bee we calculated a reinforcement index (R index) for each colour (*C_i_*) as:

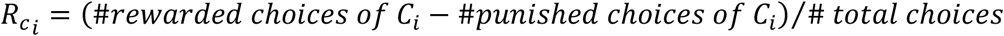

This index has a scale from -1 to +1, and colours that were experienced as punished more often than rewarded had negative R indices. Since the total number of choices both within and between trials varied between bees, the reinforcement index was normalised to the total choices bees made across all trials. Hence, the reinforcement index as defined is independent of the motivation of bees to responding to stimuli, but also shows the relative preference of bees for the different colours. We evaluated the relationship between bees’ R indices and bees’ performance in the transfer test using Spearman’s correlation test. All statistical tests were performed in MATLAB 2018 (MathWorks, Natick, MA, USA).

## Results

In the learning test bees preferred C1 (always rewarded in training) to C5 (always punished) (Fig 1c; Wilcoxon rank-sum test, z = 5.77, n = 20, p = 7.47e-10, chance level = 50%). The transfer test presented bees with a colour combination not used in training (C2 vs C4). Here bees preferred C2 to C4 (Fig 1c: Wilcoxon rank-sum test, z = 5.19, n = 20, p = 2.09e-07) demonstrating that bees preferred a stimulus with both a higher likelihood of reward and a higher ranking in a novel colour comparison.

The degree of preference differed between the learning test and transfer tests (Wilcoxon signed rank test, z = 3.92, n = 20, p = 8.79e-05). The preference for C1 in the learning test was greater than the preference for C2 in the transfer test (Fig 1c). This indicates that the degree of colour preference was not absolute (as would be expected if bees were probability maximising). Rather the degree of preference was influenced by the probability of reward and punishment received for different colours during training, which is consistent with probability matching.

There was no difference in performance in the learning, transfer and novel tests of bees that had been trained with different colour contingencies (Groups A and B, Figure S2a; Wilcoxon rank-sum test, z = -1.32, n = 10, p > 0.18). There was also no difference in performance between bees from different training protocols in the learning, transfer and novel tests (P1 and P2, Figure S2b; Wilcoxon rank-sum test, z = -1.32, n = 10, p > 0.18).

As expected, the R index differed significantly for different colours (Figure 2a,b, Wilcoxon rank-sum test, z > 3.74, n =10, p < 1.82e-4). The R index for C3 was greater than zero (Wilcoxon rank-sum test, z = 4.16, n = 10, p = 3.07e-5 for group A; z = 5.21, n =10, p = 1.82e- 7 for group A). This means that even though C3 was paired with reward and punishment in 50% of the training trials, on average bees experienced C3 as rewarded more often than punished, indicating that bees visited C3 more in rewarded trials than punished trials. In the novel test bees preferred the familiar colour (C3) more than a novel colour (Fig 1c: Wilcoxon rank-sum test, z = 5.76, n = 20, p = 7.93e-09), perhaps as a consequence of this experience in training.

**Figure 2.**
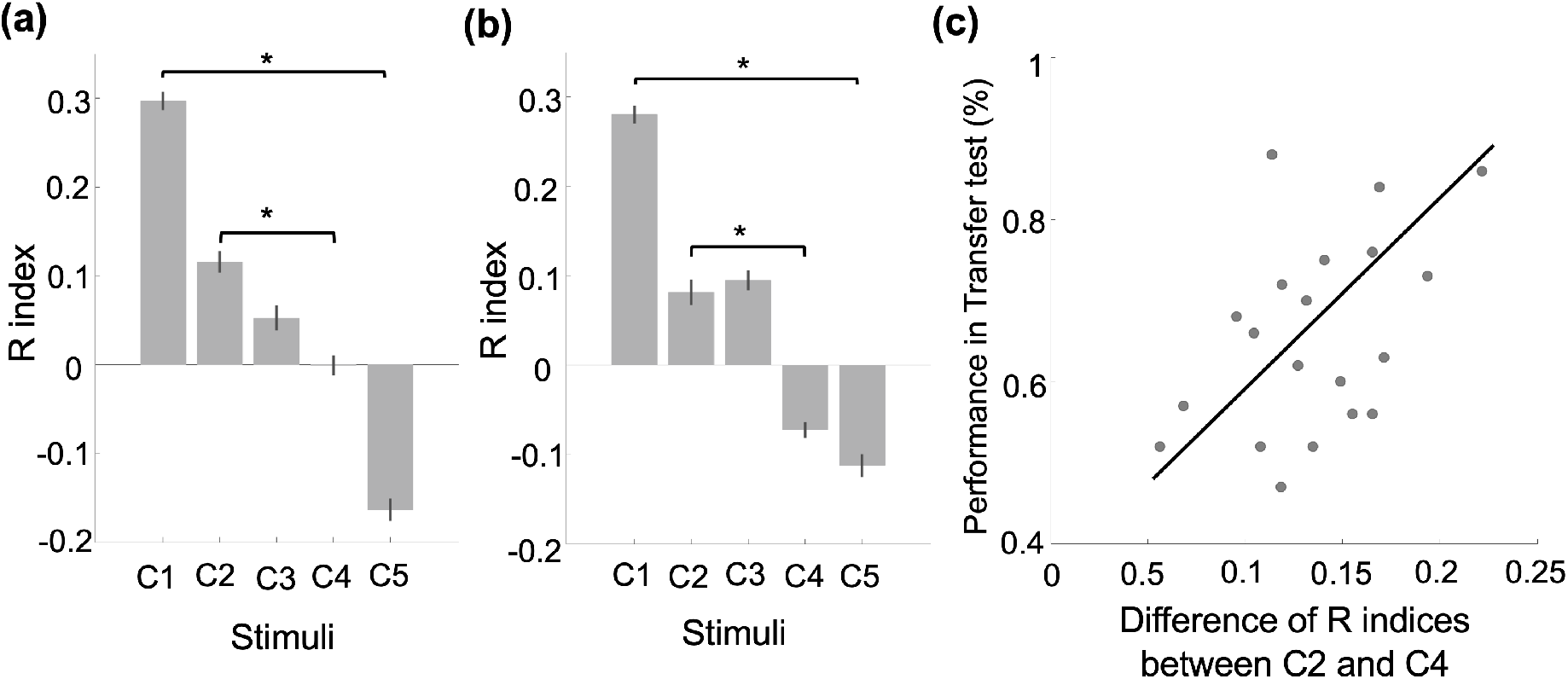
***a,b*)** Mean R index for bees for each colour (a = group A, b = group B). Bars show mean and s.e.m. of the 20 bees experience of each colour in the training phase. * indicates p<0.05 for comparing the R index of a pair of colours used in the learning test (C1 vs C5) and the transfer test (C2 vs C4). ***c)*** Relationship between performance in the transfer test and difference in R index between the two colours in the transfer test. Line indicates the linear fit to the data (Spearman’s correlation test, rho = 0.4, n = 20, p = 0.046).

Bees’ performance in the transfer test was positively correlated with the difference in their R indices for C2 and C4 (Figure 2c; Spearman’s correlation test, rho=0.4, n=20, p=0.046). Bees that experienced the greatest difference in probability of reward between C2 and C4 showed the greatest preference for C2 in the transfer test. This finding is consistent with each individuals degree of preference in the transfer test being influenced by their specific experience of reward and punishment for the two colours in training. It suggests that bees colour preferences reflect their learned history of reinforcement for each colour. It is not consistent with bees comparing and ranking colours by reward likelihood to make a choice.

### A simple neural network model sufficient for learning history of reinforcement, consistent with the neuroanatomy of the bee brain

To explore how learning of the history of reinforcement of colours in this 5-colour learning task might be achieved by honeybees we developed a neural network model inspired by the neurobiology of colour coding and colour learning in bees [20–23] (Fig 3a). Full details of the model are given in Supplementary methods. In essence our model considers how neurobiologically plausible input from different colours to the bee mushroom body might be learned if they are associated with different schedules of reinforcement.

**Figure 3.**
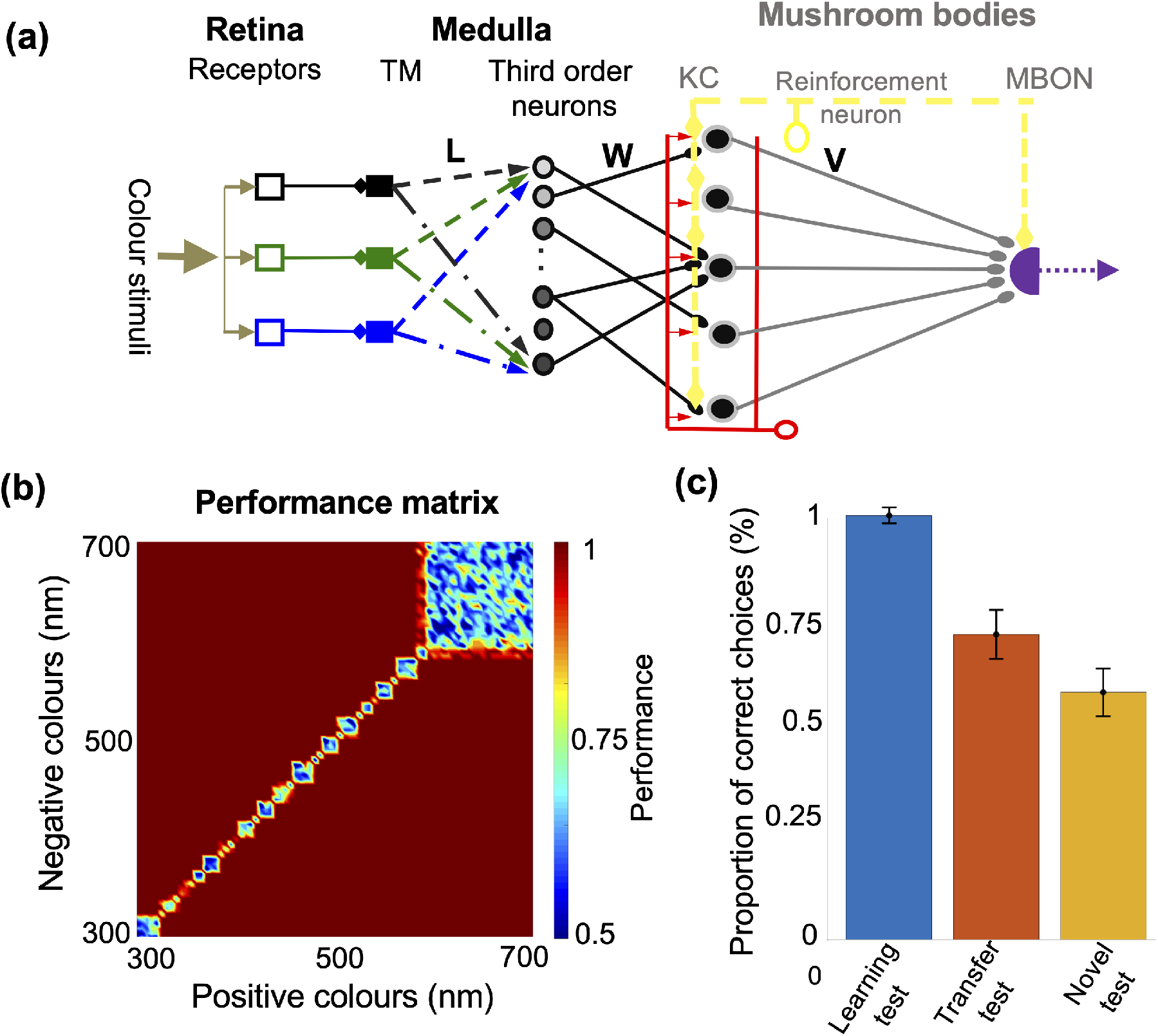
**a)** Network topology of the model of colour learning in the bee brain. Colour information is transferred from the three different types of photoreceptors to the medulla. Each receptor sends an inhibitory signal to only one type of transmedullary neuron (TM). Next, TM neurons randomly connect with the third order neurons via a connection matrix L. The vector connectivity of L was estimated from empirical data to produce a wide diversity of colour sensitive neurons. Colour information was transferred from the medulla to the Kenyon cells of the mushroom bodies (MB) via connection matrix W. Kenyon cells output to a single neuron (MBON) in the alpha lobe of the mushroom bodies via connection matrix V. The inhibitory feedback PCT pathway (red) maintains low excitability and sparse coding across the Kenyon cell population. A neuromodulatory neuron firing in response to both reward and punishment projects to both the input and output of the MB. There it acts to alter synaptic connectivity in both regions (W and V) in response to reward and punishment according to the proposed learning rule (see Supplementary Material and Method). **b)** The performance matrix of the model in a simple colour discrimination task. The colour at each matrix’s element displays the performance of the model to two monochromatic colours whose wavelengths are presented on the x and y axes. Following 10 trials with one colour associated with reward and one with punishment the model is able to discriminate all combination of colours if their wavelength distances are larger than 5 nm. **c)** Model’s performance following training with protocol P1 or P2 (Table S2). Bars were calculated from the performance of the model for 50 different initial parameters that simulated 50 different model bees that were trained to the training stimuli.

In this model, three different types of receptors, Short (S)-, medium (M)- and long (L)- wavelength-sensitive photoreceptors are stimulated by the light reflected by each colour stimulus, which we quantified as the spectral reflectance function of each stimulus (Fig. S1). Axons from colour receptors project to the medulla and make inhibitory connections with transmedullary (TM) neurons [24–27]. In bees the L-receptors project to the lamina also, [28] but in this model we consider medulla processing only.

We considered a simple circuit such that one transmedullary neuron is activated by one type of receptor only. Transmedullary neurons exhibit high spontaneous activity and receive inhibitory signals from receptors, therefore the three different types of transmedullary neurons respond to colours by decreasing their firing rate from the spontaneous rate [24]. The transmedullary neurons send either excitatory or inhibitory signals to the third order neurons (amacrine or large field neurons) in the next layer of the medulla [24, 29, 30](Fig 3a). In the model their synaptic weights (L) were estimated from empirical neurophysiological data to reproduce the diverse activity of colour sensitive neurons reported by Kien and Menzel [22, 23] (Fig S2). Previously, Vasas et al. [21] showed that in a similar model random connectivity between transmedullary neurons and third-order neurons was sufficient to describe all the documented types of colour sensitive neurons in medulla: narrow or broad-band neurons that respond to a small or large range wavelength of the light and colour opponent neurons that respond to multiple ranges of light wavelengths with either excitation or inhibition [21].

The third-order cells of the medulla (Fig 3a) project to the Kenyon cells (KCs) of the mushroom body in the collar region. W in Figure 3A describes the matrix connectivity between the third-order neurons and KCs. In the last layer of the network, Kenyon cells output to a single mushroom body output neuron (MBON) in the alpha lobe of the mushroom body *D* (*c*) through a vector of synaptic weights, *V.*

The protocerebral-calycal tract feedback pathway (Fig 3a, red) takes inputs from all Kenyon cells and sends a feedback inhibitory signal to collar region [31–35]. In the model thjis pathway is represented as a single neuron that contributes feedback gain control to the system and maintains sparse coding across the Kenyon cell population.

In the model, a single reinforcement neuron (Fig 3a, yellow) modules strengths of synaptic connectivity at both the input and output of the Kenyon cells in response to both reward and punishment. This is responsible for the changes in activity of the network during training with rewarded and punished coloured stimuli (supplementary methods, Equation 6-8).

The output of the MBON *D*(*c*) was used to evaluate the performance of the model. *D*(*c*) has a tonic firing rate, which is decreased by punishment, and increased by reward. Following training, maximal performance of the model was judged as an increase in firing rate of *D*(*c*) to maximum to a colour that had been rewarded in training, or a decrease in firing rate of *D*(*c*) to minimum to a colour that had been punished in training.

First, we assessed the performance of the model in a simple colour discrimination task. The model was presented with any pair of monochromatic colours between 300 to 700nm, one was rewarded and one punished. Prior to training the model did not differentiate between any two colours (no difference in output from *D*(*c*)). Following 10 training trials with the rewarded and punished colours the model was able to discriminate different colours in a manner consistent with performance of bees in behavioural experiments (Fig. 3B) [36, 37].

We then trained the model using protocol P1 or P2 (Table S2). Performance of the model closely matched responses of bees in the learning test (Fig 3A; compare with Fig 1C). Following training the model also successfully discriminated between a novel pair of colours and preferred the colour that had been more often associated with reward in training (Fig. 3C). Performance in this transfer test was similar to that of honey bees (Fig. 3C; compare with 1C). In both cases bees and the model matched their preference of the more rewarded stimulus to the frequency at which it had been rewarded in training.

The model did not differentiate between a novel colour and a colour that had been equally reinforced and punished during training (Fig 3C). Here performance of the model differed from performance of honey bees. Honey bees preferred a colour that had been paired with reward in 50% of trials (C3) over a novel colour (Fig 1C). But, as we noted above, bees made more visits to C3 when rewarded than when punished and hence the mean R index for C3 was positive (Fig 2a,b). In this respect training of bees differed from training in the model. When training the model the R index of a colour paired equally with punishment and reward was zero. We therefore added an additional test, and found that the model was able to successfully discriminate a colour that had been paired with reward on 66% of trials during training from a novel colour (Fig. S5, Wilcoxon rank-sum test, z = 5.31, n = 20, p = 5.62e-8).

The model allowed us to examine which elements of our network were necessary for this form of colour discrimination learning. Sparseness of colour coding in the KCs strongly affected the performance of the model: Dense coding of colours in the KC population reduced the ability to learn to discriminate colours by reward and punishment (Fig S6).

Plasticity at both the input (W, Fig 3a) and output (V, Fig 3b) of the Kenyon cells was essential for the model to correctly discriminate all colours by their history of reinforcement. In training if the weights in connection matrix W (between the third-order neurons and Kenyon cells) were fixed bees could learn to prefer a colour that was always rewarded over a colour that was always punished in training (Fig. S6b,c), but their performance in the transfer test (a comparison between one stimulus reinforced at 66% and one reinforced at 33%) was reduced when compared to the full model (Fig 3c and Fig S6b,c), and depended on both the perceptual similarity and the reinforcement history of the colours (Fig S6b,c). Plasticity in connection matrix W decorrelates the activity of KC for any two presented colours that differ in reward history, even if the population activity of third-order neurons are highly correlated (Fig S4). Hence, post training distinctive groups of KCs separately encode the colour information of each colour. This increases the ability of KCs activated by different colours to drive different levels of activity in the MBON from *D*(*c*) due to changes in connectivity resulting from different reward/penalty ratios associated with each colour during training.

If connection weights in W were fixed the degree of difference between the pattern of KC activation to two different colours correlated with the perceptual difference between the two colours (Fig S1). With this limitation the response of the model to any specific colour post training was sensitive to both the reinforcement history of that colour during training, and the reinforcement history of similar colours during training (Fig S6b,c). This effect was especially apparent in the transfer test which challenged the limited model to differentiate colours that had quite similar reinforcement histories in training (Fig S6b,c).

In the full model the plasticity rule we implemented in W was effective in decorrelating the Kenyon cell activities in response to perceptually similar colours if the colours differed in history of reinforcement during training. As a consequence, post training the full model was able to correctly differentiate pairings of colours by likelihood of reward in training only, as long as the colour wavelengths differed by more than 5 nm.

## Discussion

Our data suggest bees quickly learned a solution to a five-armed bandit task by matching their colour choices in tests to the probability of each colour being rewarded in training. Rather than comparing or ranking stimuli, bees simply aligned their likelihood of choosing a colour to their history of reinforcement with each colour. This requires nothing more than learning the properties of each stimulus, with no comparison between options. This solutions is elegant in two ways.

First, probability matching may be the ecologically optimal solution to this kind of task if reinforcement probabilities for each option are unknown or could change [1, 2, 7]. Matching choice probability to the history of reinforcement offers a simple and effective solution to the explore/exploit trade off in which an animal must optimise across exploiting a current resource type or patch, or moving to and sampling alternatives [1, 7]. Probability matching offers a solution to this classic foraging challenge without requiring a bee to compare any different types of resources. All the bee has to do is match its visits to a resource to its experience of the utility of the resource. Typical bee foraging “strategies” such as floral constancy and floral majoring and minoring can all be emergent outcomes of a simple probability matching strategy [2, 7].

Second, probability matching as a solution to a five-armed bandit task is computationally parsimonious. Niv [7] has argued that reinforcement learning is sufficient for probability matching. The mushroom body is an important brain region for learning [38–40], and octopaminergic and dopaminergic neuromodulatory neurons are essential for plastic adjustment of connection strengths in the mushroom body circuit in response to reinforcement [41–46]. Distinct but interacting dopaminergic and octopaminergic neurons encode reward and punishment in the fly brain [42], and it is likely that something similar occurs in honey bee brains [43, 44].

In our model we considered the insect mushroom body and its visual inputs (Fig. 3) as a very simple reinforcement learning system to explore the feasibility of learning to probability match in insects. In the model plasticity at both the input (calyx) and output (lobes) of the mushroom body was necessary for the model to effectively learn the history of reinforcement for different colours. Plasticity at the mushroom body input was needed for the system to be able to decorrelate the neural representations of perceptually similar colours in order to learn independent reinforcement histories for them (Fig S6). Reinforcement-related plasticity at the mushroom body calyx and lobes is feasible for insects. Hammer [47, 48] argued the modulatory neuron VUMmx1 mediates learning of sugar reward in honey bees. This neuron innovates both the calyx and lobes of the mushroom body [47]. There are other modulatory inputs (both inhibitory and excitatory) to the calyx of both bees and flies [44–46]. Strube-Bloss [39] has also argued plasticity in the calyx may be important for learning to distinguish different stimuli by reward and punishment.

Our model also emphasised the need for sparse coding of colour across the Kenyon cell population for efficient learning (Fig S5). Our mushroom body model is equivalent to a three-layer associative network. Sparse coding in the middle layer of a three-layer network is recognised to be an important feature for effective classification in such types of network [49, 50]. In honey bees sparse coding is supported by the GABAergic inhibitory feedback PCT pathway from the mushroom body outputs to the calyx [20, 44, 51-53], and such feedback is essential for complex discrimination in bees [32, 54].

Lastly we note that in this study there was no evidence honey bees ranked the colour stimuli, despite the ranking being unambiguous throughout the training. We can explain bee’s behaviour in this complex task without requiring the bees to compare the properties of any of the stimuli offered. This is perhaps a counter-intuitive way of thinking about behaviour in a choice test, but probability matching will give the appearance of choice and preference without the animal effecting any choice. This is, of course, not evidence bees or other insects *cannot* rank. As we noted above, Polistes wasps have solved a transitive inference task that controlled for reinforcement history [12] suggesting a capacity to compare and rank. Honey bees, however, failed at a similar task [55]. We note that the speed at which bees learn reinforcement history, and the effectiveness of this strategy in optimising performance in most foraging tasks may obviate the need for more complex choice strategies in most circumstances.

We also propose that given its simplicity, plausibility and ubiquity probability matching should be considered the most parsimonious explanation of behaviour in animal ‘choice’ assays, and that cognitive explanations involving overt comparison and choice should only be invoked if probability matching to reinforcement history can be categorically ruled out.

## Acknowledgements

We thank Neville Dearden from the Department of Computer Science, Sheffield University for expert beekeeping support. We thank Michael Port from Sheffield Robotics assistance in building the testing arena. We thank Vera Vasas of Queen Mary, University of London for measuring the spectra of the coloured stimuli. HM and JARM are supported by the Engineering and Physical Sciences Research Council (grant no EP/P006094/1). ABB is supported by funding by a Future Fellowship from the Australian Research Council (FT140100452), a Leverhulme Visiting Fellowship from the Leverhulme Trust and the Templeton World Charity Foundation (grant no. TWCF0266).

