## Supplemental method, figures and tables for "Honey bees solve a multi-comparison ranking task by probability matching"

### I- Supplementary Method:

#### Model of colour coding in the visual sensory system of bees

Three different types of photoreceptors (i.e. short, medium and long wavelength sensitive) are simulated by the light reflection of the stimulus. The amount of light absorbed by these photoreceptors was estimated as

$$P_m = R\int_{300}^{700} \mathrm{I}_c(\lambda)S_m(\lambda)D(\lambda)d\lambda \qquad \text{Eq. 1}$$

where $R$ describes the overall sensitivity of the receptor, $I_c(\lambda)$ is the spectral reflectance function of the stimulus, $c$, $S_m(\lambda)$ is the spectral sensitivity function of each receptor (short, medium and long), and $D(\lambda)$ is the illumination. Based on prior research on bees' photoreceptors, in this study we set perfectly uniform illumination ($D(\lambda) = 1$ at all $\lambda$) and the sensitivity factor $R = 6$.

Because of the reported non-linearity in the response of photoreceptors, the receptors' response to a colour $c$, $E_m(c)$, is calculated from the quantum catch, $E_m(c) = P_m/(P_m + 1)$.

Next, each photoreceptor sends an inhibitory signal to only one type of transmedullary cell in the medulla (TM). Hence the firing rate response of $m - th$ TM neuron is calculated with

$$r_m^{TM}(c) = r_0 + v_m E_m(c) \qquad \text{Eq. 2}$$

where $r_0$ is the high spontaneous activity of TM cells. $v_m$ is the inhibitory synaptic weights of pre-synapse TM cells which take values between [-1, 0].

The TM cells send either excitatory or inhibitory signals to the third-order cells in the next layer (Fig 3a) of the medulla through the matrix of synaptic weights $L = (L_{m,k})$. The response of the $k-th$ third-order cells is modelled as

$$r_k^A(c) = F\left(\sum_{m=1}^{3} L_{m,k} r_m^{TM}(c) + b^A;\ \alpha, \beta\right) \qquad \text{Eq. 3}$$

Where $F(x; \alpha, \beta) = A_0/\left(1 + e^{-\alpha(x-\beta)}\right)$ is the activation function and $b^A$ represents the spontaneous activity of third-order neurons. The parameters $\alpha, \beta$ control the sensitivity of the third-order neurons to the total presynaptic input. Following Vasas et al [1], the synaptic weights of the these third-order cells, $(L_{m,k})$, were fitted to the empirical data [2, 3] using a gradient descent algorithm as described in Vasas et al [1].

### Model of learning in the mushroom bodies

After fitting the third-order cells' responses to the empirical data, the firing rate of Kenyon cells (KC) of the mushroom bodies (Fig 3a) is modelled as:

$$r_p^{KC}(c) = F\left(\sum_{k=1}^{22} W_{p,k} r_k^A(c) - I_f;\ \alpha, \beta\right) \qquad \text{Eq. 4}$$

where $W_{p,k}$ is the synaptic connectivity between $k-th$ third order neurons and $p-th$ KC in the mushroom bodies. $I_{f=}\ \sum_{m=1}^{N_{kc}} Q_m . r_m^{KC}(c)$ is the input of the inhibitory feedback pathway to the KC that is obtained from the average activity of Kenyon cells. The synaptic weights $Q_m$ control the sparseness of activity of the KC. Higher values of $Q_m$ increase the inhibitory input to the KCs (Eq. 4). This results in reduced or no activity in some KC. Hence, the population activity of KC becomes more sparse. Setting $Q_m$ to 0 effectively models the system without any inhibitory feedback to the KC, which produces dense activity in the KC population.

The outputs of all KC stimulate a single mushroom bodes output neuron, $D(c)$. Through a vector of synaptic weights, $V = (v_1, v_2, \ldots v_i);\ i = 1{:}N_{kc}$, The activity of this neuron in response to the colour, $c$, is expressed by

$$r^D(c) = F\left(\sum_{m=1}^{N_{kc}} V_m . r_m^{KC}(c);\ \alpha, \beta\right) \qquad \text{Eq. 5}$$

For simplicity, we set the same activation function, $F(x; a, b)$ for all firing rate models of third-order cells, KC and the decision neuron (Eqs. 3, 4 and 5) with fixed parameters: $A_0 = 100$ spike/ sec and $\alpha = 0.05$ and $\beta = 50$.

We propose that the difference in responses of $D(c)$ to the rewarded and punished stimuli must be increased during the training phase. Hence, the optimal synaptic weights, $V^{opt}$ and $W^{opt}$, could be obtained from maximizing the cost function that represents the difference between the activity of $D(c)$ to all pairs of the positive ($c_p$) and negative stimuli ($c_n$).

$$\langle W, V\rangle^{opt} = \underset{\langle C_p, C_n\rangle}{argMax}\ [r^D(c_p) - r^D(c_n)] \qquad \text{Eq. 6}$$

If we assume $E = r^D(c_p) - r^D(c_n)$ in the equation 6, as the cost function, we have

$$\frac{\partial E}{\partial V_i} = r_i^{KC}(c_p) f\left(\sum_{m=1}^{N_{kc}} V_m r_m^{KC}(c_p); a, b\right) - r_i^{KC}(c_n) f\left(\sum_{m=1}^{N_{kc}} V_m r_m^{KC}(c_n); a, b\right) \qquad \text{Eq. 7}$$

and

$$\frac{\partial E}{\partial W_{i,j}} = V_j\, r_i^A(c_p) f\left(\sum_{k=1}^{22} W_{p,k,} r_k^A(c_p) - I_f; a, b\right) f\left(\sum_{m=1}^{N_{kc}} V_m r_m^{KC}(c_p); a, b\right) - V_j\, r_i^A(c_n) f\left(\sum_{k=1}^{22} W_{p,k,} r_k^A(c_n) - I_f\,; a, b\right) f\left(\sum_{m=1}^{N_{kc}} V_m r_m^{KC}(c_n)\,; a, b\right) \quad \text{Eq.8}$$

where $f(.)$ Is the derivation of the activation function $F$.

Finally, the reinforcement neuron (Fig 3a) makes a reward- or punishment- modulated connection with $D(c)$ and KC. $\delta(t)$ presents the reinforcement signal that depends on whether a stimulus is paired with reward or punishment ($\delta(t) = 1$), or with no stimulus ($\delta(t) = 0$). Hence, the synaptic weights of the last two layers, $V$ and $W$ might be increased or decreased from stimuli presented at step $t$ to $t + 1$ according to the following learning rules:

$$V_i^{t+1} = V_i^t + \eta \frac{\partial E}{\partial V_j}\, \delta(t) \quad \text{Eq. 9}$$

and

$$W_{i,j}^{t+1} = W_{i,j}^t + \eta \frac{\partial E}{\partial W_{i,j}}\, \delta(t) \quad \text{Eq. 10}$$

where $\eta$ is the time constant that controls that learning rate at which the weights change.

### II- Supplementary Figures

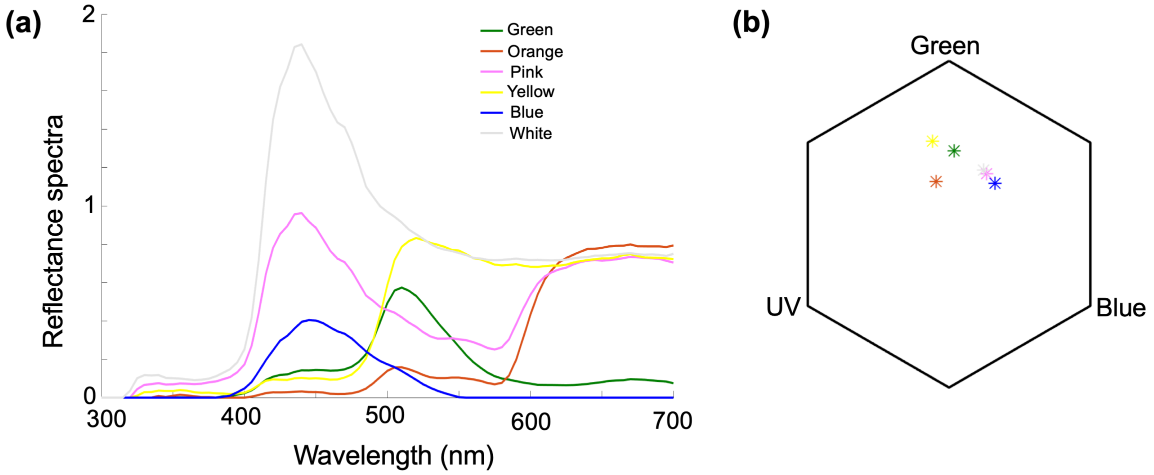


**Figure S1. Colour spectrum measurements of stimuli used in this study**. ***a)*** Relative spectral reflectance plot of each of the colours used in the training phase and the novel tests. The curves show the spectral refection of stimuli used in training phase and the tests. The colour of curves represents the colour of stimuli. ***b)*** Loci of colours in bee colour space, describing the range of colours a bee can see given their three photoreceptors maximally sensitive to UV, Blue, and Green light. The colored dots indicate each of the stimulus colours used in the training and tests.

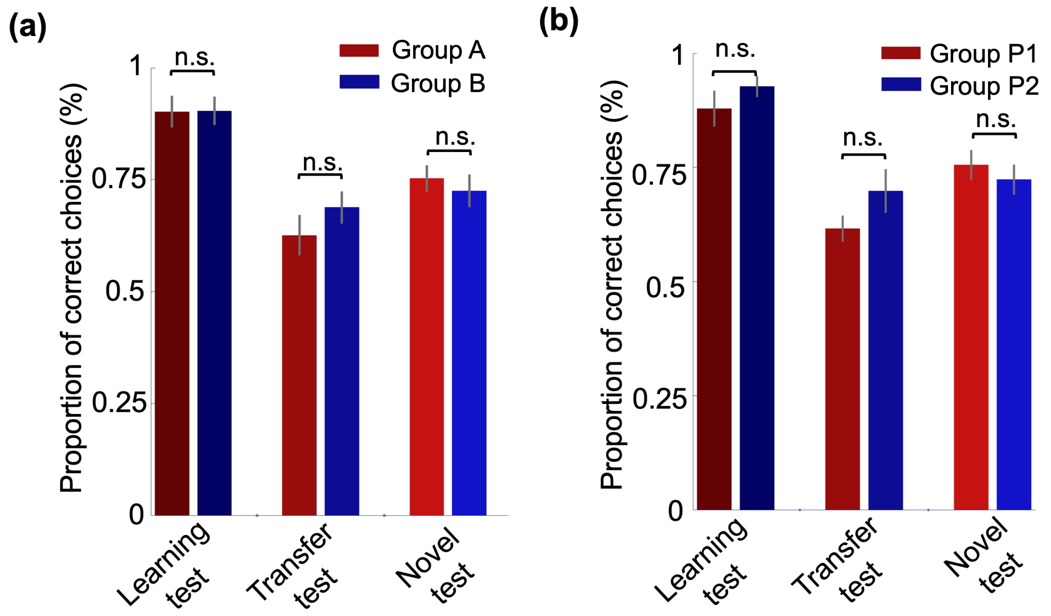


**Figure S2**. Bees' performance in the tests. There was no significant difference between ***(a)*** groups of bees trained with different colours (A vs B) and ***(b)*** groups of bees trained with different order of colour presentation (P1 vs P2).

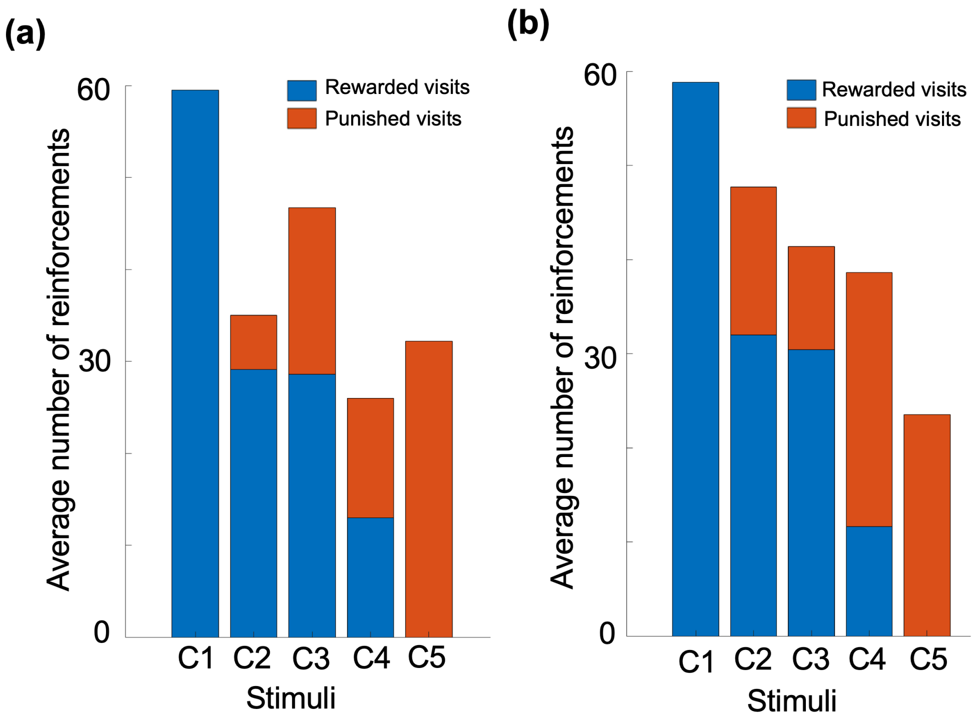


**Figure S3.** Number of visits to stimuli that were reinforced and punished in the training trials**.** The stacked bars show the average number of punished and rewarded visits that bees experienced from each colour in the training phase (a) Group A; b) Group B).

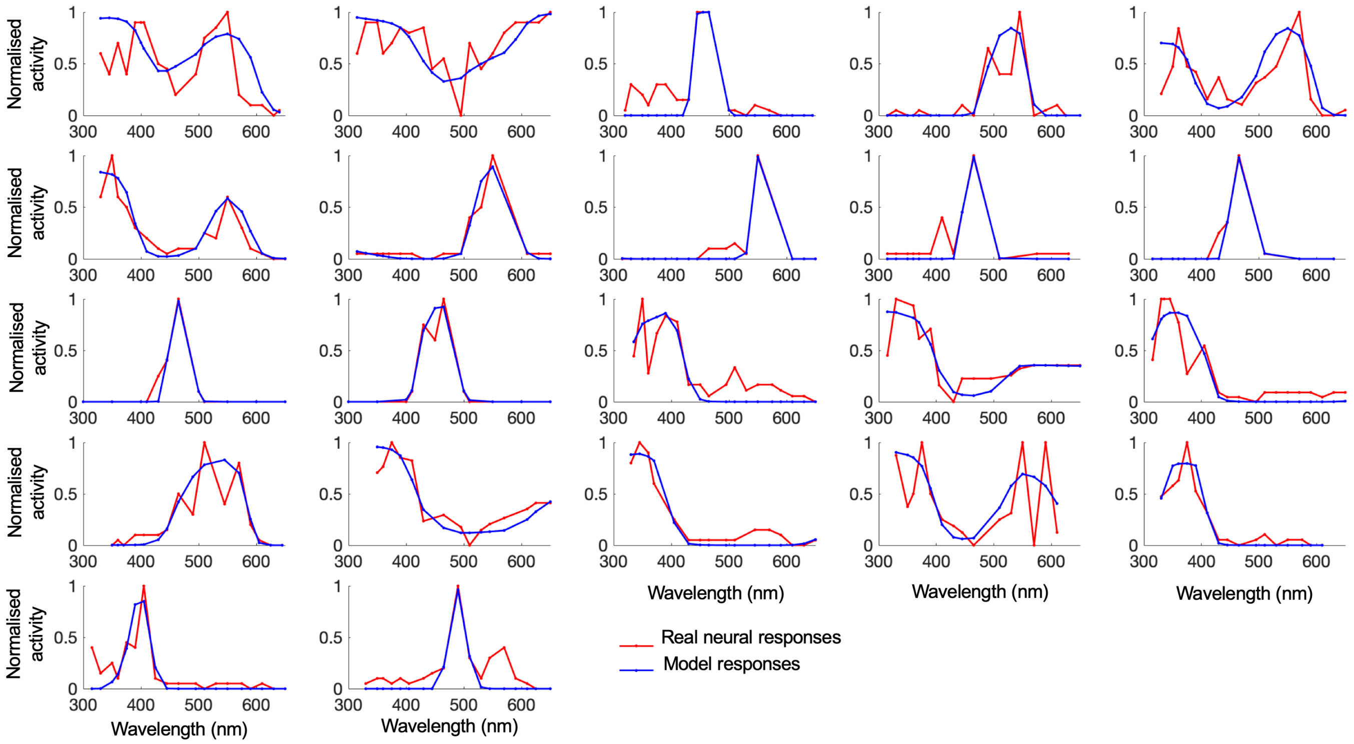


**Figure S4: A diverse activities of medulla neuron generated by the model.** Plots show the spectral tuning curves of third-order neurons. The red curves show the empirically measured spectral tuning curves reported in {Kien, 1977 #14645;Kien, 1977 #14695}. The blue curves show the model responses of the third-order neurons in the medulla to different colours.

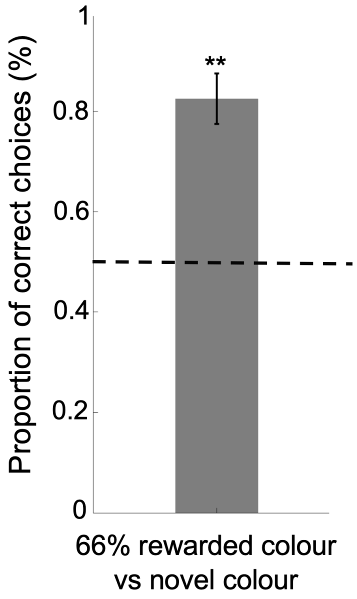


**Figure S5: Model's performance in discrimination 66% rewarded colour from the novel colour.** The bar graph shows the Mean ± SE of the model's performance in discriminating a colour rewarded 66% of time in training from a novel colour. Data generated from 50 different initial parameters that simulated 50 different model bees following training with protocol P1 or P2 (Table S2).

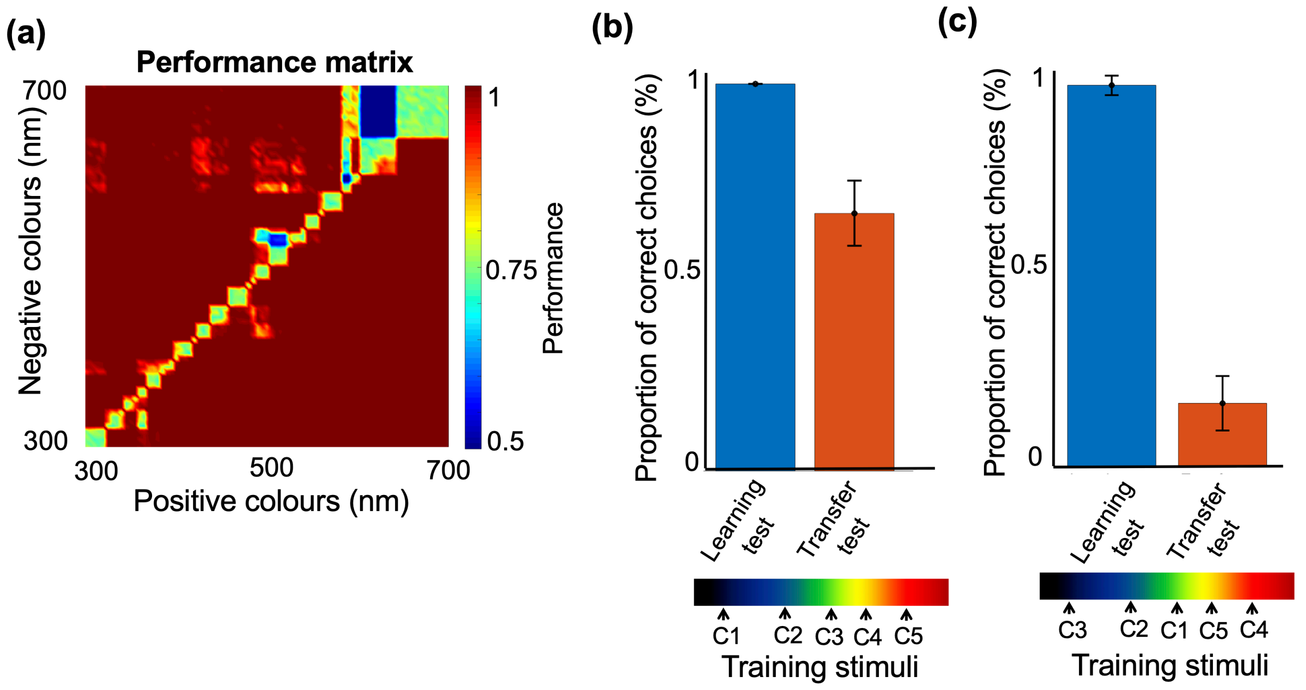


**Figure S6: a)** The performance of the model with low sparseness in the Kenyon cell population. The matrix shows the performance of the model in the colour discrimination task. The colour of each matrix element displays the performance of the model to two monochromatic colours whose wavelengths are presented on the x and y axes. The performance of the model in colour discrimination is reduced when the number of KCs activated by any colour is increased (compare with Fig. 3b). **b & c)**. Performance of the model with connection plasticity in layer V only (Fig 3). The model was trained in the history of reinforcement paradigm with two different colour sets, shown beneath panels b and c. This model with no plasticity in connection matrix W (Fig. 3) is not able to reproduce the performance of bees in the transfer test (compare with Figs. 1C and 3C). The performance of the model in the transfer test is sensitive to the set of training colors. The model chose a 66% rewarded colour more than a 66% punished colour if the five training colours were ordered from from low wavelength to high wavelength by reinforcement history during training (b). However the model did not learn to prefer the 66% rewarded colour if training colours were randomly assigned a reinforcement history (c). Bars were calculated from the performance of the model for 50 different initial parameters each initial parameter provides the responses of a model bee.

### III- Supplementary Tables

| Stimulus | % trials rewarded | % trials punished | Colour Pairs |
|---|---|---|---|
| C1 | 100 | 0 | C1 vs C2<br>C1 vs C3<br>C1 vs C4<br>C1 vs C5 |
| C2 | 66 | 33 | C2 vs C1<br>C2 vs C3<br>C2 vs C5 |
| C3 | 50 | 50 | C3 vs C1<br>C3 vs C2<br>C3 vs C4<br>C3 vs C5 |
| C4 | 33 | 66 | C4 vs C1<br>C4 vs C3<br>C4 vs C5 |
| C5 | 0 | 100 | C5 vs C1<br>C5 vs C2<br>C5 vs C3<br>C5 vs C4 |
| (Novel colour) | 0 | 0 | … |

**Table S1:** Summary of training trials**.** Each colour was paired with all others, except for pairing C4 with C2. This pairing was excluded from the training procedure to use as a novel pair in the transfer test. In each pairing the higher ranked colour (red) was rewarded and the lower ranked colour punished (blue).

**Protocol P1**

| # Bouts | Stimuli at each bout |
|---|---|
| 1 | C1 vs C2 |
| 2 | C3 vs C5 |
| 3 | C1 vs C4 |
| 4 | C2 vs C5 |
| 5 | C3 vs C4 |
| 6 | C1 vs C3 |
| 7 | C4 vs C5 |
| 8 | C2 vs C3 |
| 9 | C1 vs C5 |
| 10 | C1 vs C2 |
| 11 | C3 vs C5 |
| 12 | C1 vs C4 |
| 13 | C2 vs C5 |
| 14 | C3 vs C4 |
| 15 | C1 vs C3 |
| 16 | C4 vs C5 |
| 17 | C2 vs C3 |
| 18 | C1 vs C5 |

**Protocol P2**

| # Bouts | Stimuli at each bout |
|---|---|
| 1 | C3 vs C5 |
| 2 | C1 vs C2 |
| 3 | C1 vs C4 |
| 4 | C2 vs C5 |
| 5 | C1 vs C3 |
| 6 | C3 vs C4 |
| 7 | C1 vs C5 |
| 8 | C4 vs C5 |
| 9 | C2 vs C3 |
| 10 | C1 vs C5 |
| 11 | C3 vs C4 |
| 12 | C2 vs C3 |
| 13 | C4 vs C5 |
| 14 | C1 vs C3 |
| 15 | C2 vs C5 |
| 16 | C1 vs C4 |
| 17 | C1 vs C2 |
| 18 | C3 vs C5 |

**Table S2 & 3: Training sequences.** Two different protocols for training were used. Each protocol was created by ensuring bees experienced each pair of colours in each half of the 18 training bouts. Otherwise colour pairings were ordered randomly. 10 bees were trained with P1, and 10 with P2.

### IV- Supplementary Video

**Video S1:** Sample video of honey bee in transfer test.
